# Bacterial lipopolysaccharide reduces the stability of avian and human influenza viruses

**DOI:** 10.1101/146266

**Authors:** Christopher Bandoro, Jonathan A. Runstadler

## Abstract

Commensal bacteria can promote or reduce the severity of viral infection and disease progression in their hosts depending on the specific viral pathogen^1^. Influenza A virus (IAV) has a broad host-range, comprises many subtypes, and utilizes different routes of transmission including the fecal-oral route in wild birds^2^. It has been previously demonstrated that commensal bacteria can interact with the host’s immune system to protect against IAV pathogenesis^3,4^. However, it is unclear whether bacteria and their products may be interacting directly with IAV to impact virion stability. Herein we show that gastrointestinal (GI) tract bacterial isolates in an *in vitro* system significantly reduce the thermal stability of IAV. Moreover, bacterial lipopolysaccharide (LPS), found on the exterior surfaces of bacteria, was sufficient to significantly decrease the stability of both human and avian viral strains at the physiological temperatures of their respective hosts, as well as in the aquatic environment. Subtype and host-origin of the viruses were shown to affect the extent to which IAV was susceptible to LPS. Furthermore, using a receptor-binding assay and transmission electron microscopy, we observed that LPS binds to and affects the morphology of influenza virions.

---

Influenza A virus (IAV) is a global threat, infecting 5-10% of adults and 20-30% of children globally every year^5^, and infections in humans and highly-pathogenic IAV outbreaks in livestock substantially burden the economy^6,7^. All past pandemics of IAV in humans and outbreaks in livestock have origins in viruses that previously circulated among wild aquatic birds, which are the natural reservoir for the virus^8^. In wild birds, the virus transmits primarily via the fecal-oral route^9^. After infecting the gastrointestinal (GI) tract, virions shed in feces contaminate aquatic habitats facilitating transmission to new hosts. In all these environments, virions encounter upwards of 10^11^ bacteria per milliliter or gram^10,11^.

Commensal bacteria and their products can indirectly protect against IAV infection by interacting with the host’s immune system (Fig. 1A). In the absence of commensal bacteria, mice produced impaired type I/II interferon responses, CD4/CD8 T cell responses, and antibody production to IAV infection^3,4^. Moreover, mice pretreated with bacterial lipopolysaccharide (LPS), a product present on the exterior surface and shed by all Gram-negative bacteria, triggered a TLR4 mediated antiviral response to protect the hosts from lethal infection with IAV^12,13^. In contrast, LPS binds directly to the capsid protein of poliovirus, increasing cell attachment and the ability of the virions to remain infectious at increased temperatures^14^, and LPS binding to mouse mammary tumor virus (MMTV) enables increased immune evasion and successful transmission of the virus^15,16^. In the case of influenza, it is unclear whether commensal bacteria and LPS are interacting directly with IAV in addition to their indirect effects on the immune system.

**Figure 1:**
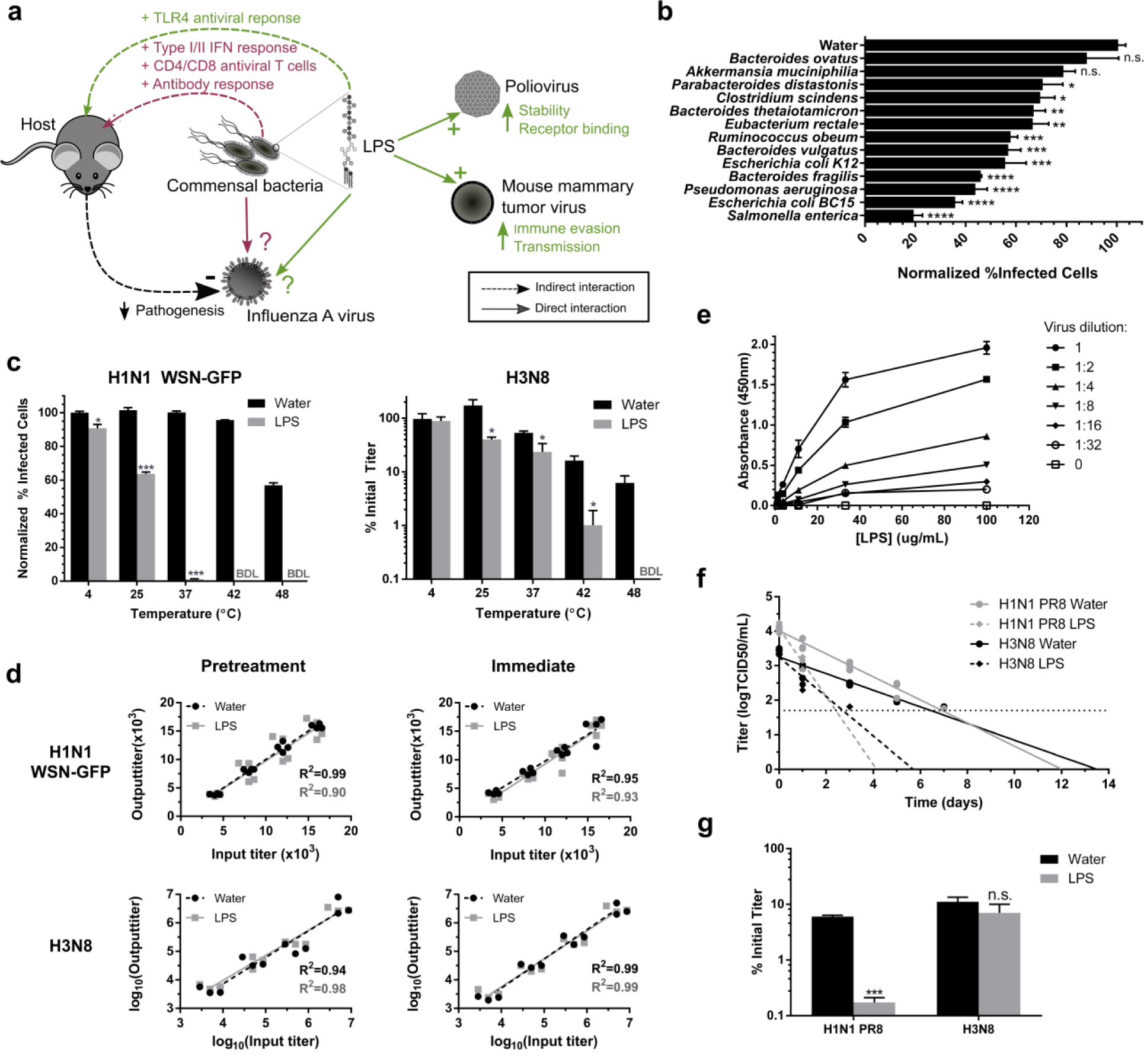
Commensal bacteria and lipopolysaccharide directly reduce the stability of influenza A virus. **a,** Schematic summarizing the indirect and direct effects of bacteria and lipopolysaccharide (LPS) on influenza A virus (IAV) and other enteric viral pathogens. b, Stability of a human H1N1 WSN-GFP virus after incubation for 1 h at 48 °C with heat-killed gastrointestinal tract bacterial isolates normalized to a water control (N=4). BDL = below detectable limit. c, Stability of a human H1N1 WSN-GFP and an avian H3N8 after incubation for 1 h at various temperatures with LPS (N=3). d, Output titers of H1N1 WSN-GFP and H3N8, across a range of input titers, from tissue-culture cells either 1) pretreated with LPS or water control and then infected, or 2) infected immediately after mixing the viruses with LPS or water (N=4). e, Binding of biotinylated-LPS to H1N1 PR8 after incubation for 1 h at 37 °C (N=2). f, Long-term persistence of a human H1N1 PR8 and avian H3N8 in water (N=13, 12), or water containing diluted LPS (N=6, 7) at 25 °C. The horizontal dashed line represents the limit of detection. g, Aquatic freeze-thaw stability of a human H1N1 PR8 and avian H3N8 in LPS or water control (N=4). Data are represented either as individual replicates or as the as mean ± SEM. Statistical significance was assessed using a, a Dunnett’s multiple comparison test, c,g, a student’s t-test, or d,f, an F-test on the linear regressions., ***p < 0.001, ** < 0.01, * <0.05, n.s. non-significant.

We first tested whether GI tract-derived bacteria can affect the stability of IAV. A panel of thirteen heat-killed bacteria derived from the GI microbiome (Table S1), standardized by protein content, and a water control were incubated individually with a human WSN H1N1-GFP virus for 1h at 48 °C. Stability was assessed by measuring the percentage of infected cells (those expressing GFP) by flow cytometry after an overnight infection. Eleven of the thirteen bacterial strains significantly decreased the stability of the virus after incubation compared to the water control (Fig. 1B). Our results indicate the reduction in infectivity was not due to the diluted bacterial products being cytotoxic (Fig. S1A), nor was the decrease in infectivity due to the presence of bacteria or their products limiting the susceptibility of host-cells to infection (Fig. S1B). This suggests that the observed reduction in thermal stability may be due to the virus interacting directly with the bacterial cells and/or their products. Because there was a wide range in the extent to which bacterial strains reduced viral stability, the specific composition of a host’s microbiome could influence its susceptibility to infection by IAV. A recent study found that the cloacal microbiome of wild mallards infected with IAV differed from those who were IAV-negative, specifically in the *Firmicutes*, *Proteobacteria* and *Bacteroidetes* phyla^17^. Although it is challenging to determine whether these differences in the microbiome preceded or were the result of IAV infection, bacteria representing these phyla used in our study all reduced IAV stability.

Given that bacterial LPS is present in milligrams/milliliter concentrations in the animal GI tract^18^, and that it interacts directly with poliovirus^14,19^ and mouse mammary tumor virus (MMTV)^16,20^, we hypothesized that LPS may be interacting with IAV and affecting its stability. Incubating a human WSN H1N1-GFP and an avian H3N8 subtype influenza with 1 mg/mL of *E. coli* O111:B4 LPS for 1 h significantly reduced the stability of both viruses in a temperature-dependent manner (Fig. 1C). Intriguingly, the stability of the human and avian viruses were respectively reduced by 82-fold and 16-fold in LPS compared to water (p <0.001 and 0.01) at 37 °C and 42 °C, the physiological temperatures of their respective hosts. Furthermore, with temperature held constant (37 °C), LPS reduced the stability of IAV in a concentration dependent manner (Fig. S2). We used our *in vitro* system to determine whether the observed reductions in infectivity in the thermal stability assays were due to LPS interacting with the virus directly or through the indirect response of infected cells. Cells were either pretreated with LPS prior to infection or not and then were treated concurrently with a mixture of virus and LPS. Across a range of virus titers tested, the addition of LPS did not reduce the expected titer in our system for both scenarios (Fig. 1D), suggesting that the reduction in infectivity observed was due to LPS interacting directly with the virus, rather than the tissue-culture cells mounting an antiviral response. This was further supported by the observation that detoxified LPS, a modified version of LPS lacking the immunogenic lipid A portion of the molecule, also reduced viral stability (Fig. S3). We also observed that biotinylated-LPS binds to H1N1 PR8 virions in a solid-phase receptor binding assay (Fig. 1E). With 2.0x10^7^ virions, the dissociation constant (Kd) was 24 µg/mL of LPS (R^2^=0.99). Since the HA glycoprotein of IAV binds to sialylated receptors on host cells, we postulated that LPS binding is mediated through HA. However, pretreating the virus with antibodies against HA or its sialic acid receptors failed to block binding to LPS (Fig. S4). LPS could be interacting with virions in an HA-independent manner, either through the viral neuraminidase or potentially host-proteins derived from the host cell membrane as with MMTV^16^.

Aquatic habitats serve as long-term reservoirs for avian IAVs and are important to sustain transmission of the viruses in wild birds^21,22^. Abiotic factors, including temperature, pH, and salinity, that affect the persistence of IAV in water are well established ^23^, but it is unclear how biotic factors, including bacteria and their products, affect virus stability. Avian H3N8 and human H1N1 PR8 viruses persisted significantly longer in water compared to water containing 100 µg/mL of LPS at 25 °C (respective p= 0.002 and <0.002) (Fig. 1F). Using the slopes of the regression analyses, for each treatment we calculated the log10 reduction time (Rt), defined as the length of time for the viral titer to decrease by 90%^24^. LPS reduced the Rt of the avian virus compared to water by 58%, and to a greater extent for the human H1N1 by 66%. Viruses circulating in the aquatic environment must also be able to withstand the natural freeze-thaw cycles that rapidly inactivate the virions^25^. We found that LPS significantly reduces the freeze-thaw stability of the human H1N1 PR8 virus by 34-fold compared to water (p< 0.001), whereas the avian H3N8 virus was not significantly affected (Fig. 1G). Bacterial products such as LPS may be reducing persistence and stability of IAV in the environment, which has as significant implications for modeling the emergence of novel strains from animal reservoir hosts.

Our long-term aquatic persistence results suggested that the avian isolate may be more resistant to LPS than the human virus tested. In addition, it has been reported that the microbiome content of IAV infected mallards was significantly affected by hemagglutinin (HA) subtype of the virus ^17^. To examine the extent to which the reduction in stability may be conserved across viruses we compared the thermal stabilities in *E. coli* LPS and water at 48 °C for 1 h of a panel of viruses that represented i) phylogenetically-distinct HA subtypes H1, H3, H4, H5, and H12 (Fig. 2A) and ii) different host-origins (avian, human, seal). All of the strains tested had significantly reduced stability in LPS compared to water (Fig. 2B), suggesting that LPS appears to have a universal effect of reducing the thermal stability of influenza viruses. Measurements of relative stability of each virus in LPS compared to water (LPS:water) did not follow a normal distribution (Shapiro-Wilks test, p<0.05) and were tested with a non-parametric Kruskall-Wallace H test. A significant difference was detected between subtypes (p<0.001), as well as host origin (p<0.001). The subtypes H4, followed by H5 and H12 (all avian origin) showed the highest stability in LPS, compared to H1 and H3 (Fig. 2C). We further analyzed H1 and H3 subtypes because for these two subtypes measurements of LPS:water stability were made for two or more hosts. For H1, virus originating from avian hosts was significantly more stable in LPS, compared to human viruses (DF=1, p=0.006). For H3, avian and seal viruses were more stable in LPS compared to human viruses, though not at a statistically significant level (DF=2, p=0.055).

**Figure 2:**
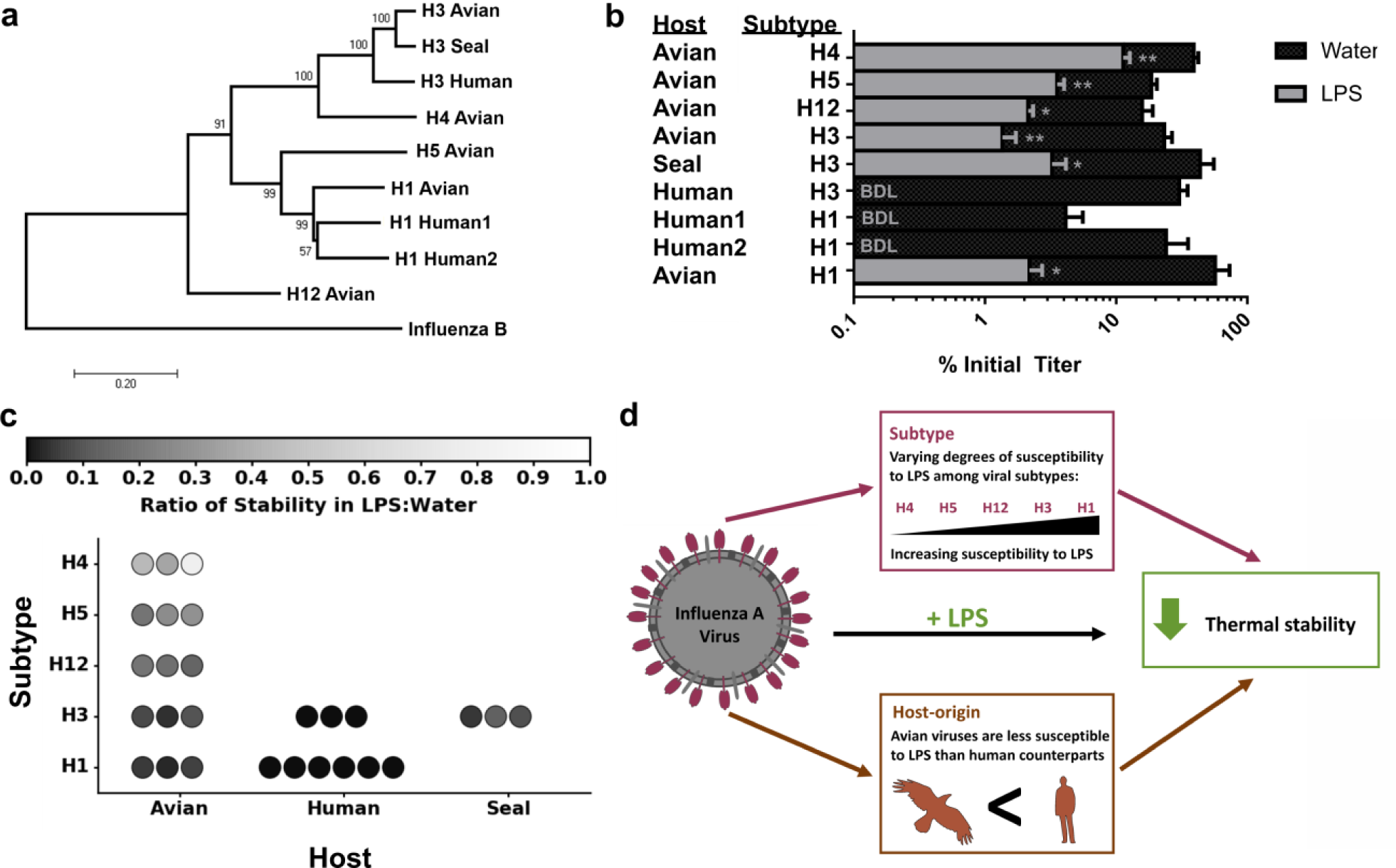
Host-origin and subtype affect the susceptibility of influenza A virus to lipopolysaccharide (LPS). **a,** Phylogenetic tree constructed using the hemagglutinin (HA) gene of nine influenza A viruses with influenza B virus included as an outgroup. Bootstrap percentages for the internal nodes are displayed. Scale bar shows the evolutionary distance (number of base substitutions per site). b, Stabilities of these viruses after incubation for 1 h at 48 °C in lipopolysaccharide (LPS) or water control (N=3). Data represented as mean ± SEM. Statistical significance was assessed using a student’s t-tests to compare LPS and water **p-value <0.01, * <0.01. BDL = below detectable limit of 0.025% Initial Titer. c, Ratio of stability in LPS:Water of the viruses across Hosts and Subtypes. Each replicate is represented as a circle. d, Schematic representing the summary of the results.

Our data suggest that subtype and host-origin of IAV influence the ability of the virus to withstand inactivation (Fig. 2D). Most notably, avian-derived strains of IAV were more stable in LPS compared to the human-derived stains. Given avian IAV transmits primarily via the fecal-oral route ^9^, compared to the respiratory transmission of human IAV^26^, and that the bacterial load in the GI tract is higher than airways ^27,28^, avian strains may be adapted to the high concentrations of LPS that they encounter inside the host and in feces.

Because LPS affects IAV in a temperature dependent manner via direct binding, reducing the stability of the virus at increased temperatures, we next hypothesized that LPS may be compromising the structural integrity of the virions’ envelopes. H1N1 PR8 virions incubated in water at 37 °C for 1 h retained their spherical morphology by transmission electron microscopy (TEM), whereas virions incubated with LPS had envelopes that were more noticeably deformed (Fig. 3A). To quantify whether there was a change in virion morphology, we measured the length:width ratio of the virions (Fig. S5). LPS treated virions had significantly different length:width ratio distribution compared to control virions using a Kolmogorov-Smirnov (KS) test (p <0.001) (Fig. 3B), but did not result in increased virion aggregation (Fig. 3C). Therefore LPS causes an increase in the proportion of virions with envelope deformations, which may account for the decrease in stability observed.

**Figure 3:**
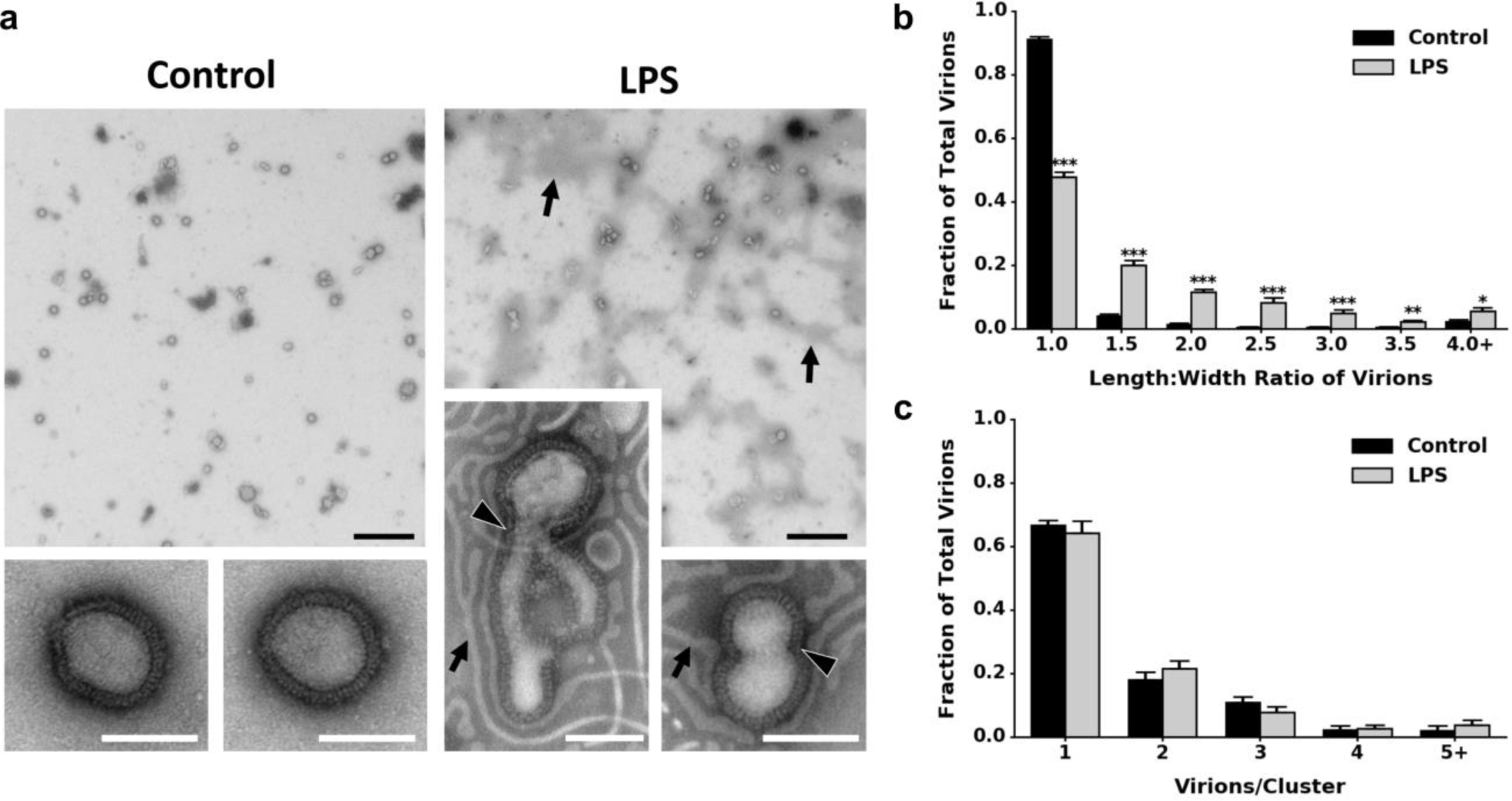
Bacterial lipopolysaccharide (LPS) comprises viral envelope integrity but does not does not affect aggregation. **a,** Transmission electron micrographs of H1N1 PR8 after incubation for 1 h at 37 °C in either water (control) or lipopolysaccharide (LPS). Representative micrographs were taken at 4,800x and 49,000x magnifications. LPS ribbons (arrows) and compromised virion membranes (arrowhead) are labeled. Scale bars: black, 1 μm; white, 100 nm. b-c, Fractions of total virions in water control (N=559) and LPS (N=579) binned by b, the length:width ratio of the virions to assess differences in viral envelope morphology, or c, the number of virions per viral cluster to assess differences in virion aggregation. Fractions of virions in were calculated based on the total number of virions in each micrograph. Data are represented as the mean ± SEM of 9-10 micrographs. Statistical significance was assessed by a student’s t-test. ***p-value <0.001, ** <0.01, * <0.05.

In conclusion, our study provides new evidence that LPS directly affects the stability of IAV by binding directly to and altering the morphology of influenza virions, suggesting that bacteria within hosts and in the external environment can limit the transmission of influenza virus. Our results are presented mainly in the context of IAV transmission within its natural avian reservoir, however we found that human-derived viruses could be more susceptible to bacterial products. Understanding the extent to which bacteria interact with and affect the infectivity of IAV may suggest novel pathways to the development of therapeutics to prevent or treat respiratory infections. Moreover, the overuse of antibiotics in poultry and swine, in addition to generating and disseminating antibiotic resistant strains of bacteria^29^, may be increasing their risk of IAV infection. However, our results stand in contrast to those found with poliovirus^14,19^ leading us to propose that interaction with LPS is highly dependent on the virion structure. Additional studies monitoring the effects of antibiotics and changes in the bacterial microbiome on the susceptibility of host animals, as well as humans, to pathogenic viruses will help clarify the extent to which bacterial-viral interactions may be influencing the severity and epidemiology of viral infection.

## Methods

**Bacterial strains, viruses and cell-lines:** Cultures of GI tract and fecal microbiome bacterial isolates from humans and mice (see Table S1) were heat-killed at 95 °C for 10 min and standardized by their protein content using a bicinchoninic acid (BCA) assay (ThermoFisher). Viruses used in this study are listed in Table S2. We also used a reverse-genetics engineered H1N1 WSN PB1flank-eGFP (obtained from the laboratory of Jesse Bloom), referred to in this study referred to as H1N1 WSN-GFP, in which the PB1 gene’s coding sequence was replaced by a gene encoding enhanced green fluorescent protein (eGFP) ^30^. Viruses were propagated in either Madin Darby Canine Kidney (MDCK)cells (obtained from American Type Culture), MDCK-SIAT1-CMV-PB1 cells (obtained from the laboratory of Jesse Bloom), or embryonated chicken eggs (obtained from Charles River Laboratories) for 72 h. The avian H3N8 and human H1N1 PR8 were further purified and concentrated through a 30% sucrose cushion during ultracentrifugation ^31^.

**Ethics Statement.** Cultures of GI tract and fecal microbiome bacterial isolates were obtained from an existing collection of isolates from the laboratory of Timothy Lu. The sources of the bacterial isolates from humans were anonymized. Embryonated chicken eggs, obtained from a vendor (Charles River Laboratories), were infected on days 9 to 11 during development as reviewed and approved by the Committee on Animal Care at the Massachusetts Institute of Technology (Protocol Number: E15-02-0218).

**Growth and Infection Media.** MDCK cells and MDCK-SIAT1-CMV-PB1 cells were grown in DMEM (Hyclone) supplemented with 10% FBS (Seradigm), 100 U/mL penicillin and 100 µg/mL streptomycin (Sigma) at 37 °C with 5% CO_2_. H1N1 WSN PB1flank-eGFP infections were carried out in MDCK-SIAT1-CMV-PB1 cells in OptiMEM (Hyclone) supplemented with 0.01% FBS, 0.3% BSA, 100 U/ml penicillin and 100 ug/ml streptomycin, and 100 ug/ml calcium chloride at 37 °C with 5% CO2. Infections involving all other the viruses were carried out in DMEM supplemented with 0.2% BSA (ThermoFisher), 25mM HEPES (Corning), 100 U/mL penicillin and 100 µg/mL streptomycin at 37 °C with 5% CO2.

**Virus titers and infectivity.** H1N1 WSN PB1flank-eGFP was initially titered as described previously ^30^. Briefly, serial dilutions of the stock virus were allowed to infect 1.0×10^5^ MDCK-SIAT1-CMV-PB1 cells for 16 h at 37 °C with 5% CO2. The % infected cells was determined by flow cytometry (Accuri C6, Accuri Cytometers), using uninfected control cells as the baseline and setting 0.5% of the control cells as GFP-positive (FlowJo). We then used the Poisson equation to calculate the number of infectious units (IU)/mL of the virus in initial stock. For the stability assays, viruses were added to cells as described above along with a standard curve of known quantities of virus in parallel to ensure they were in the linear range of the assay. To compare samples across independent replicates, we normalized the % infected cells to that of the water control. All other viruses were titered in MDCK cells by tissue-culture infectious dose 50 (TCID50) assays. Briefly, 3.0×10^4^ MDCK cells were seeded into 96-well plates (VWR) overnight and then viruses were serially diluted across the plate, incubated for 2 h to allow attachment, washed, and then returned to incubate for 72 h at 37 °C with 5% CO2. Presence or absence of cytopathic effect (CPE) was observed and the TCID50/mL was calculated using the Reed-Muench method ^32^.

**Thermal stability assays:** To test whether bacterial products may be affecting the thermal stability of influenza virus, 1.0×10^6^ infectious units (IU)/mL of H1N1-GFP was mixed into water or the heat-killed bacterial isolates, standardized to 1 mg/mL protein in water, and incubated at 48 °C for 1 h. Similarly, to test the effects of LPS on influenza stability, 1.0×10^6^ IU/mL of H1N1 WSN-GFP or 5.0×10^6^ TCID50/mL of the other viruses (Table S2) were spiked into either water or 1 mg/mL of purified *E.coli* O111:B4 LPS (Sigma) at a range of temperatures for 1 h. Stability for H1N1 WSN-GFP was measured by determining the %Infected Cells, normalized to water, by flow cytometry (see Supplemental Experimental Procedures). For all other viruses, we calculated the %Initial Titer remaining after incubation by TCID50 assay ^33^ as a measure of stability. We also compared the stability of the avian H3N8 and human H1N1 WSN-GFP at 42° and 37 °C, representing internal temperatures of their respective hosts, in purified LPS from *E. coli* O111:B4, *Salmonella enterica* serotype minnesota (Sigma), and *Pseudomonas aeruginosa* 10 (Sigma) and water. For all thermal stability trials student t-tests were applied to determine whether there was significant difference between the virions in water and LPS. To compare the thermal stabilities of the panel of IAV strains tested, we calculated the ratio of their stability in LPS:Water (%Initial Titer in LPS/%Initial Titer in Water).

**Thermal stability controls:** As a control to determine whether the change in infectivity was caused by bacteria or LPS affecting the susceptibility of the MDCK or MDCK-SIAT1-CMV-PB1 cells, we examined the output titers of a range of known input titers of avian H3N8 and human H1N1 WSN-GFP. Cells were pretreated for 2h at 37 °C (the length of time the cells were incubated with viruses + LPS in the assay before being washed) with diluted heat-killed *S. enterica* or *E. coli* LPS prior to infection, or were infected immediately with mixtures of the viruses and 1 mg/mL of *S. enterica* or LPS. An F-test was performed on linear regressions of the samples pretreated or immediately infected with LPS and water to test whether the slopes of the lines were significantly different. A WST-1 assay (G-Biosciences) was used to assess the cytotoxicity of the bacteria to the cells without the presence of virus. As an additional control, virions were also incubated with 1 mg/mL of detoxified *E. coli* O111:B4 LPS (Sigma) which is missing the immunogenic lipid A portion.

**Aquatic environmental stability:** For the long-term persistence experiment, 1.0×10^4^ TCID50/mL of avian H3N8 or human H1N1 PR8 were added to water or water with 100 µg/mL of LPS and incubated at 25 °C. Viruses were periodically removed and titered immediately over 7 days to determine the quantity of infectious virions remaining. Data points were log10 transformed and then fit with a linear regression using GraphPad to calculate the log10 reduction time (Rt) which is the time required for infectivity to decrease by 90% (or 1 log10 TCID50/mL) ^24,34^. An F-test was performed to compare the slopes of the linear regressions.

Freeze-thaw stability was tested by adding 1.0×10^6^ TCID50/mL of human H1N1 PR8 or avian H3N8 virus to either water or 1 mg/mL of *E. coli* LPS. Samples were frozen at −20 °C for 3-5 days and then thawed at 25 ° C. Stability was assessed similar to the thermal stability experiments by measuring the titer prior to freezing and after thawing the viruses.

**Phylogenetic analysis:** A panel of viruses (see Table S2) was selected based on diversity of hemagglutinin (HA) subtypes and host-origin of the viruses. The nucleotide sequences of the viruses’ Segment 4 gene, which encodes HA, were used to construct a phylogenetic tree. We used MEGA7 ^35^ to compute the evolutionary distances via maximum composite likelihood method ^36^ and assessed the evolutionary history using the Neighbor-Joining method ^37^. Influenza B virus B/Durban/39/98 was included as outgroup in the analysis.

**Binding:** To test whether LPS binds to virions, we used a modified version of a solid-phase influenza receptor binding assay described previously^8^. Briefly, H1N1 PR8 virions, at varying dilutions, were bound to fetuin A (Sigma) coated plates overnight at 4 °C. After washing, dilutions of biotinylated *E. coli* O111:B4 LPS (Invivogen) were added incubated at 37 °C for 1 h. LPS binding was detected using HRP-streptavidin(ThermoFisher), TMB substrate (Thermofisher), and 0.2 M sulfuric acid. Absorbance was measured at 450 nm. Binding Kd was calculated using a non-linear regression with GraphPad software. To assess whether LPS binding was mediated by HA, virions were pretreated for 1 h at room temperature prior to LPS with dilutions of monoclonal antibodies against the globular HA head (anti-HA head, provided by Peter Palese’s lab), polyclonal antibodies against H1N1 (anti-H1N1, Thermofisher), or with α2,6 and α2,3 sialyllactose (Carbosynth).

**Transmission electron microscopy (TEM):** 1.0×10^8^ TCID50/mL of H1N1 PR8 virions were mixed with 1 mg/mL of *E. coli* LPS or water control and then incubated at 37 °C for 1 h. Carbon/formvar film supported Cu grids were glow discharged and then placed on a drop of influenza virus mixtures and then stained with 1% (wt/vol) phosphotungstic acid. Samples were examined using a Tecnai G2 Spirit BioTWIN Transmission Electron Microscope and representative TEM images were taken at 4,800× and 49,000× magnifications. To test whether LPS was affecting the shape of the virions, the length and the perpendicular bisecting width of the virions were measured using ImageJ at 4800× magnification across micrographs for each treatment. The ratio of length: width of each virion was calculated. An example can be seen in S3 Fig̤ To determine whether LPS was affecting the aggregation of virions, virions in different viral clusters (cluster sizes of 1, 2, 3 etc.) were enumerated as above in each treatment. Fractions of virions in each length:width ratio bin or cluster were calculated and normalized based on the total number of virions in each micrograph.

**Statistical analyses.** Statistical analyses were performed in Graphpad Prism 6 software, JMP Pro 12, and Python v2.7 with Numpy v1.12 and Scipy v0.18. Student’s t-tests were performed to compare the means between two groups (parametric). To compare multiple groups, we assessed normality via a Shapiro-Wilks test, and then computed a one-way ANOVA followed by a post-hoc Dunnett’s multiple comparison test (parametric) or Kruskall-Wallace H test (non-parametric). To compare distributions we used the non-parametric Kolmogorov–Smirnov (KS) test to test the null hypothesis that the distributions were the same. Statistical significance threshold was assessed at p-values < 0.05. Refer to the Fig. legends or tables to find the statistical tests, exact values of n, what n represents, the definition of center, dispersion and precision measurements.

**Data availability.** New sequence data for four avian hemagglutinin genes used in the phylogenetic analysis are available on Genbank: A/mallard/Interior Alaska/12ML00957(H1N1)/2012 (KY750593), A/mallard/Interior Alaska/12ML01123(H5N2)/2012 (KY750601), A/mallard/Interior Alaska/12ML00831(H4N6)/2012 (KY750585), A/mallard/Interior Alaska/12ML00678(H12N5)/2012 (KY750577).

## Acknowledgments

The authors acknowledge funding support for this work from the National Institute of Allergy and Infectious Diseases (NIAID) Centers of Excellence for Influenza Research and Surveillance (CEIRS) (HHSN272201400008C), MIT’s Center for Environmental Health Sciences (CEHS) (P30-ES002109), and the National Science and Engineering Research Council of Canada (NSERC). We thank Justin Bandoro and Nichola Hill for help with statistical analyses and revising; Mark Mimee & Rob Citorik, Jesse Bloom, and Wenqian He for providing bacterial isolates, the WSN-GFP virus system, and antibodies respectively; Jamie Cheah at the Koch Institute at MIT for help with ultracentrifugation; Marcus Parrish and Bob Croy at MIT for help with flow cytometry; Maria Ericsson at the Harvard Medical School Electron Microscopy Facility for their help with transmission electron microscopy; Kimberly Davis for sequencing the mallard IAV isolates used in this study; and all the Runstadler Lab members for their valuable critique and input.

## Supplementary information

Supplementary Tables 1-2 and Supplementary Figures 1-5 can be found within the Supplementary Information PDF.

